# Evolutionary rates for multivariate traits: the role of selection and genetic variation

**DOI:** 10.1101/002683

**Authors:** William Pitchers, Jason B. Wolf, Tom Tregenza, John Hunt, Ian Dworkin

## Abstract

A fundamental question in evolutionary biology is the relative importance of selection and genetic architecture in determining evolutionary rates. Adaptive evolution can be described by the multivariate breeders’ equation 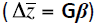, which predicts evolutionary change for a suite of phenotypic traits 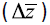 as a product of directional selection acting on them (***β***) and the genetic variance-covariance matrix for those traits (**G**). Despite being empirically challenging to estimate, there are enough published estimates of **G** and ***β*** to allow for synthesis of general patterns across species. We use published estimates to test the hypotheses that there are systematic differences in the rate of evolution among trait types, and that these differences are in part due to genetic architecture. We find evidence some evidence that sexually selected traits exhibit faster rates of evolution compared to life-history or morphological traits. This difference does not appear to be related to stronger selection on sexually selected traits. Using numerous proposed approaches to quantifying the shape, size and structure of **G** we examine how these parameters relate to one another, and how they vary among taxonomic and trait groupings. Despite considerable variation, they do not explain the observed differences in evolutionary rates.

## Introduction

Predicting the rate and direction of phenotypic evolution remains a fundamental challenge in evolutionary biology [1–4]. Empirical studies have demonstrated that most traits are heritable [5–8] and can respond to selection – a prediction borne out by an abundance of short- (e.g. [9–11] and long-term (e.g. [9,12–14] artificial selection experiments targeting single traits. However, in most biological systems, the targets of selection are suites of traits. Furthermore, different traits are tied together by genetic associations (typically quantified as covariances), and consequently selection on one trait can lead to evolutionary changes in other traits [7,8,11,15–21]. Indeed, genetic covariation between traits appears to be ubiquitous and has the potential to shape the evolution of associated traits [7,10,17,18,20,22,23]. Therefore, to improve our understanding of phenotypic evolution it is necessary to invoke a multivariate perspective [5,17–19,24].

The evolutionary response of a suite of traits can be predicted by the multivariate breeder’s equation 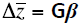 where 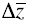 is the vector of responses in phenotypic means for the suite of traits, **G** is the additive genetic variance-covariance matrix and ***β*** is the vector of linear (directional) selection gradients [5–8]. The importance of **G** to phenotypic evolution can be illustrated using the concept of “genetic degrees of freedom” [9,11,15]. Whenever there is genetic covariance between them, the number of trait “combinations” in **G** that can respond to selection can be considerably smaller than the actual number of measured traits. This can be true even when each trait in **G** is heritable and all pairwise genetic correlations between them are less than one [1–3,9,11,25]. This reduced dimensionality constrains the population to evolve in a genetic space with fewer dimensions than the number of traits (and trait combinations) potentially under selection. A matrix whose variance is concentrated in one or a few dimensions can exhibit “lines of least evolutionary resistance” (LLER); directions in which the multivariate evolutionary response can proceed more rapidly than in others [15]. The presence of these LLERs can have a major influence in biasing the direction of evolutionary trajectories (Figure 1; ref.s [7,11,15–20]), making the **G** matrix more informative about the short term capacity of a population to respond to selection (i.e. its evolvability) than the heritabilities of individual traits [7,10,17,18,20,22,23].

**Figure 1.**
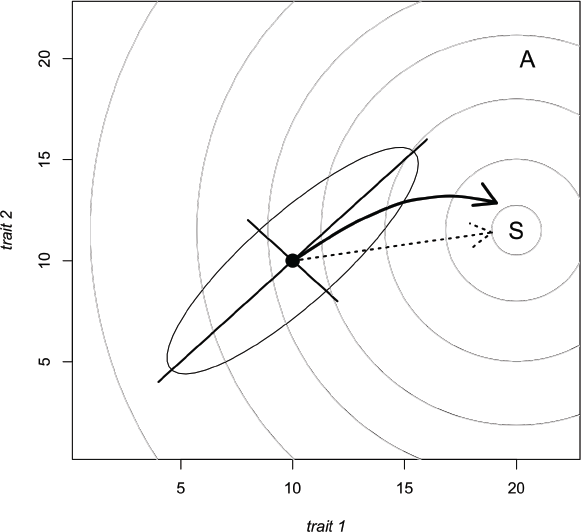
The effect of **g**_max_ on the response to selection where traits genetically covary. The axes represent the breeding values for 2 hypothetical traits. The population mean is at the solid point and the surrounding ellipse is the 95% confidence region for the distribution of trait values about the mean. That these traits covary is evident as the ellipse is at an angle relative to the trait axes. The axes of the ellipse represent the 2 orthogonal directions (eigenvectors) of variance present – there is more standing genetic variance along the major axis (**g**_max_) than the minor axis. They grey lines are ‘contours’ on a fitness landscape, with an adaptive peak at ‘S’. Rather than evolving directly toward the peak (dashed arrow), the influence of **g**_max_ may cause the population to evolve along an indirect course (bold arrow). In some cases this may even, result in the population evolving toward an alternate fitness peak (e.g. at ‘A’, modified contours not shown) in line with **g**_max_, even though it is more distant from the current mean.

A variety of measures have been proposed as proxies for the evolutionary potential of a population. Most current approaches represent a function of the components of the multivariate breeder’s equation: **G**, ***β*** and 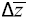 [5,17–19,21,24]. Unfortunately, few studies simultaneously estimate more than one of these components. The notable exceptions suggest that the structure of **G** plays an important role in directing phenotypic evolution [26–29]. Even fewer studies provide direct estimates of observed rates of evolution [30,31]. However, many individual estimates of selection and evolutionary rates exist in the literature and evolutionary research has benefitted from reviews that synthesize these parameters [30–38]. There is considerable variation in the strength of selection across different trait types and fitness measures [33,34,38], as well as over time (but see ref.s [36,39,40]. On average, linear selection appears stronger on morphological than life-history traits and both linear and quadratic selection is stronger when acting on mating success and fecundity compared to viability [1–4,33,38]. However inferences from such studies are subject to methodological debate [5–8,35] and potentially publication biases [9–11,40]. In particular, there has been disagreement about trait scaling, and how it influences estimates and broader evolutionary conclusions [19,22,41].

Although they have not received the same attention as selection gradients, reviews based on published genetic parameters show clear differences across trait types. Morphological traits generally have higher heritabilities than life-history traits, with physiological and behavioural traits intermediate between these extremes [9,12–14,32], but see [6–8,11,15–21]. Sexual traits have also been shown to have higher additive genetic variances compared to non-sexually selected traits [7,10,17,18,20,22,23,42], although this finding is based on few studies. As discussed above, trait scaling has been shown to alter the observed patterns [19,22,41].

There have been even fewer attempts at synthesis from a multivariate perspective. Notably, Kirkpatrick [20], Kirkpatrick & Lofsvold [9], Agrawal & Stinchcombe [23], and Schluter [11,15] collected small samples of **G** matrices from the literature and found that much of the available variance was concentrated in the first few dimensions. This suggests that few genetic degrees of freedom may be the norm, but we know of no systematic review that reveals how general this pattern is or whether it differs across taxa or trait types. Likewise, although reviews on the rate of contemporary microevolution suggest that rapid evolution should be viewed as the norm rather than the exception [15,30,31], a comprehensive review of evolutionary rates across different taxa and trait types does not currently exist.

We compiled a database of reported genetic parameters from the literature to ask whether different types of traits evolve at different rates, and whether such differences correlate with differences in selection, in patterns of genetic (co)variation or both. We performed a quantitative literature review, to examine whether observed rates of evolutionary response differ across trait types (morphological, life-history and sexual) in plants and animals. We relate these observed rates of evolutionary response to estimates of linear and quadratic selection, as well as measures that capture the size, shape and structure of **G** [7,11,15–20], to determine whether there is an association across trait types and taxa. We find some evidence that sexual traits evolve faster than other traits in animals but not in plants, where life-history traits evolve fastest. These increased rates of evolution do not appear to be attributable to the same cause however. In plants we find that selection also appears to be strongest on life-history traits, whereas in animals selection on sexually selected traits appears to be stronger than on life-history but indistinguishable from that on morphology. We then examined how the measures used to capture the size, shape and structure of **G** vary among trait types and between taxa, but find that this incompletely explains the observed pattern of evolutionary rates. In addition, we compare the various measures based upon **G**, and show that for these empirically observed matrices, many strongly co-vary.

## Methods

All data and scripts containing our analyses can be downloaded either from DRYAD (doi:xxx) or github (https://github.com/DworkinLab/Pitchers_PTRS2014).

### Compilation of Database

We compiled our datasets by searching for publications on the ISI Web of Science database between March 2006 and August 2012. We then refined this preliminary list of references on the basis of their title, abstract and keywords and attempted to obtain the full text for all papers included in the dataset.

Rates of evolution have been measured using a number of different units, most prominently darwins [7,10,17,18,20,22,23,43,44] and haldanes [5,17–19,21,24,43,45]. Measurements in darwins have proved most appropriate for researchers studying evolution on macro-evolutionary scales (e.g. paleontologists), since they express the rate of evolution per million years (although there are known methodological issues with making comparisons [44,46]). However, for our purposes rates expressed in haldanes are the appropriate unit as they measure change per generation and are used to measure evolution on a micro-evolutionary scale – the scale over which **G** may be important. We therefore compiled a database of evolutionary rate measured in haldanes *only*. We performed searches for the terms ‘*rate of evolution’*, ‘*rate of adaptation’*, ‘*haldane*s’, ‘*response to selection*’ and ‘*experimental evolution*’. This process was aided considerably by making use of the measurements from the studies previously compiled by Hendry *et al* [26–29,47]. Where studies reported the results of experimental evolution without explicitly reporting a rate of response, we contacted the authors to ask for the data needed (e.g. generation time) to calculate a rate in haldanes, standardizing traits as necessary. Previous work has shown that even with log transformation of ratio scale data (where means and variances might co-vary), this had little influence on overall estimates for haldanes [31].

For the database of selection gradients, we began with the database compiled by Kingsolver *et al* [30,31,33,37], and supplemented this with additional measures from work published after 2001 by searching for the terms ‘*natural selection’*, ‘*sexual selection*’, ‘*selection gradient*’ or ‘*selection differential*’. Unlike Kingsolver *et al* [30–38] we included both field and laboratory studies. While there has been discussion about the effects of trait scaling (mean vs. standard deviation) on estimates of selection [19,35], we have only included estimates standardized using the approach as advocated by Lande and Arnold [21], as this has been most broadly used.

For the **G** matrix dataset we searched the Web of Science database using the terms ‘*G-matrix*’ (or ‘*G-matrix*’), ‘*covariance matrix*’ (or ‘*co-variance matrix*’ or ‘*(co)variance matrix*’) or ‘*quantitative genetics*’. We recorded **G** matrices expressed both as genetic (co)variances (provided we were able to mean-standardize them, following [19]) and as genetic correlations and narrow sense heritabilities. Where possible (i.e. where estimates of phenotypic variance had been presented alongside genetic correlations and heritabilities) we back-calculated the genetic variances and covariances as: 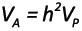 and 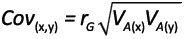 where *V_A_* and *V_P_* are the additive genetic and phenotypic variances, *h^2^* is the narrow sense heritability and *r*_G_ is the genetic correlation between traits *x* and *y*. In cases where matrices were incomplete we contacted the author(s) to request the missing estimates. We thus have two **G** datasets; correlation matrices and covariance matrices. Since we found correlations to be reported more often than covariances, the correlation dataset is a superset of matrices that includes those in the covariance dataset. Trait scaling for the co-variance matrices is discussed below. In a number of cases matrices had component traits that had been measured in difficult-to-compare units (e.g. both a length and a volume), or where traits were expressed as residuals (e.g. from regression against size). In these cases we excluded these from the reported analysis, but inclusion had little effect on the results. A number of matrices were also found to include cells with correlations >1 and in these cases we excluded the offending matrix.

### Defining Trait Categories and Measures

Since we wished to make comparisons across different ‘trait types’ (*sensu* [33,34,38]), it was necessary to assign our measurements from the literature into categories. We chose three trait categories: life-history, morphological and sexually selected traits. It is relatively straightforward to separate life-history from morphological traits and the majority of measurements in the literature fall into these two categories. In animals, we defined sexual traits as those where we were able to find at least one study demonstrating the trait was subject to female preference or used in male-male competition. For plants, we defined floral morphology as sexually selected [36,39,40,48]. Thus, for both plants and animals, our sexually selected and morphology categories are not mutually exclusive. In an attempt to reduce error in our study, traits that did not fit clearly into one of our three categories were excluded from our dataset. For **G** matrices whose component traits did not all fit the same category, we split the matrix to produce sub-matrices relating to traits only within a single category. Where matrices contained a single trait whose category differed from all others in the matrix we removed that trait from the matrix.

When making comparisons across our trait categories, we acknowledge that our classifications may not be directly equivalent in plants and animals. We therefore included a ‘taxon’ category in our statistical models. The list of individual measures of evolutionary rate was treated as a single response variable, as were the standardized selection gradients.

In our analysis of the **G** data, we wished to capture those attributes of **G** that might be expected to influence the rate of evolutionary change. Matrices vary principally in terms of size and structure. While numerous studies suggest that the alignment of axes of **G** with ***β*** is likely to be important, the nature of the data we were able to compile does not allow us to quantify alignment. Instead (as outlined below) we utilized a number of scalar measures derived from **G**, meant to capture aspects of the size and structure as a means to express evolutionary potential, All of the measures we used are summarized in Table 1. One general concern is that not all of the measured we used explicitly accounted for the number of traits included in the matrix (i.e. *n_D_*). While, in general the number of traits seemed to have a small influence on these measures (Figures 4 & 5), we also took several steps to account for these effects, such as including number of traits as a linear co-variate in the models (below) and also by examining the effects of scaling *n_D_* by either trait number or its square (“effective subspace”, as suggested by one of the manuscript referees). In none of these cases did it substantially alter the results. While we use the name “effective dimensionality” for *n_D_*, as proposed by Kirkpatrick [20], this measure actually captures aspects of matrix eccentricity, not dimensionality.

**Table 1.**
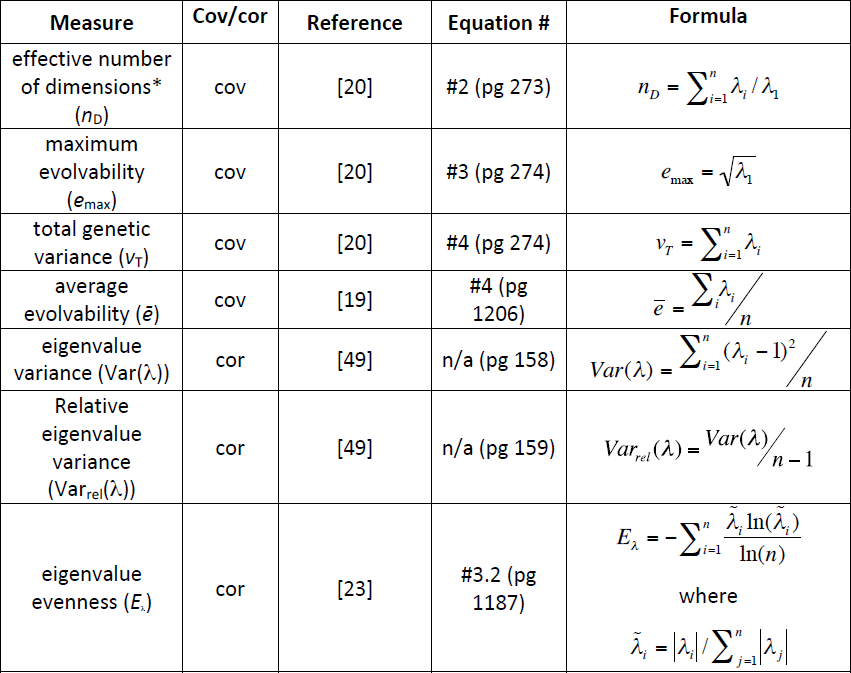
**G**-matrix measures used in this study. Eigenvalue variance, relative eigenvalue variance and eigenvalue evenness are calculated from correlation matrices, whereas the other four metrics are calculated from covariance matrices. In all formulae λ are eigenvalues and *n* is the number of traits in the matrix.* *n_D_* does not measure dimensionality per se, but eccentricity.

For the dataset of **G** as mean-standardized covariance matrices we used the three **G**-structure measures suggested by Kirkpatrick [20]: ‘total genetic variance’ (*tgv*), ‘maximum evolvability’ (*e*_max_) & ‘effective number of dimensions’ (*n_D_*), and also Hansen and Houle’s [19] ‘average evolvability’ (*ē*). For the dataset of correlation matrices, we calculated Pavlicev *et al*.’s [49] eigenvalue variance (Var(λ)) and relative eigenvalue variance (Var_rel_(λ)) and also Agrawal & Stinchcombe’s [23] eigenvalue evenness (*E_λ_*). Both sets of **G** matrix measures are defined in Table 1.

While we present results from analyses of both the (co)variance and correlation matrix datasets, it is important to note that results are not directly comparable between them, since it is well known that different methods of scaling (i.e. mean-standardizing (co)variance matrices vs. effectively variance-standardized correlation matrices) produce fundamentally different results for genetic attributes [6,19,35]. Furthermore, though the correlation matrix dataset is larger, we note that the covariance – not correlation – matrix is the current standard expression of **G** used for response to selection [21], and rates calculated from correlation matrices would also not be directly comparable to those calculated from covariance matrices.

### Statistical Analyses

Analyses were performed using **R** (version 2.13.0; ref. [50]); we fit generalized linear mixed-effect models using the MCMCglmm package (version 2.15; ref. [51]). A large proportion of studies reporting selection gradients also reported standard errors or confidence intervals (from which standard errors can be calculated). As noted by Kingsolver *et al* [38], this allows for the application of formal meta-analyses, and we followed their lead in modelling selection data with a meta-analysis including random-effects to account for study- and species-level autocorrelation. We analysed estimates of standardised selection gradients (***β***) expressed as absolute values.

We found that standard errors or confidence intervals were reported much less frequently among studies of **G** or rates of evolution, and so we were unable to account uncertainty in the estimates of **G** in these analyses as we had for selection, though the model structure we used was otherwise similar. We fit a set of models, and then evaluated model fit by comparing Deviance Information Criterion values (DIC) [52], and confirmed our selections by refitting the model set using reduced maximum likelihood (lme4 package [53]) and comparing fits using Akaike and Bayesian Information Criterion scores (AIC/BIC) and likelihood ratio tests using a parametric bootstrap. The selected models for each dataset are described in Table 2, and full model sets are available with the data and scripts on Dryad and github. Since we modelled the magnitude (absolute value) of our response variables, we used the folded normal distribution [38]. We therefore extracted the posterior distributions of solutions, took the mean and standard deviation from these distributions and applied these to the folded normal distribution. We then report the mean and credible intervals from these corrected distributions [38].

**Table 2.** The main effects included in the final models for each analysis. (Effects of ‘trait type’ refer to life-history, morphology or sexual and ‘taxa’ to plant or animal. ‘Study type’ refers to field observation or experimental evolution. Random effects of ‘study’ and ‘species’ refer to models where an intercept was fitted to each species and study, and the random effect of ‘trait type:species’ indicates where both a species-level intercept and a species-level trait type effect were fitted.) Full sets of models can be found in the scripts and data on Dryad.

| measure | fixed effects | random effects |
| --- | --- | --- |
| Rate (Animals) | trait type + study type | study + trait type:species |
| Rate (Plants) | trait type | species |
| $\vert \beta\vert $ | trait type + taxon + trait type x taxon | study + species |
| (G) $nD$ | trait type + taxon + trait no. | study |
| (G) $e_{\max}$ | trait type + taxon + trait no. | study + trait type:species |
| (G) $tg\nu$ | trait type + taxon + trait no. | study + trait type:species |
| (G) $\bar{e}$ | trait type + taxon + trait no. | study + trait type:species |
| (G) $\text{Var}(\lambda)$ | trait type + taxon + trait no. | study |
| (G) $\text{Var}_{\text{rel}}(\lambda)$ | trait type + taxon + trait no. | study |
| (G) $E_{\lambda}$ | trait type + taxon + trait no. | study |

In total we used 2571 estimates of the rate of evolutionary response (measured in haldanes); there were comparatively few estimates for plants, with no estimates available on the observed rate of evolution for sexually selected (floral) traits. This imbalance caused our estimates to be unstable so we modelled plant and animal rates separately. We had 776 estimates of ***β*,** but **G** is reported less frequently in the literature (Table 3) and our sample size of **G** measures was 81 (co)variance matrices and 221 correlation matrices.

**Table 3.**
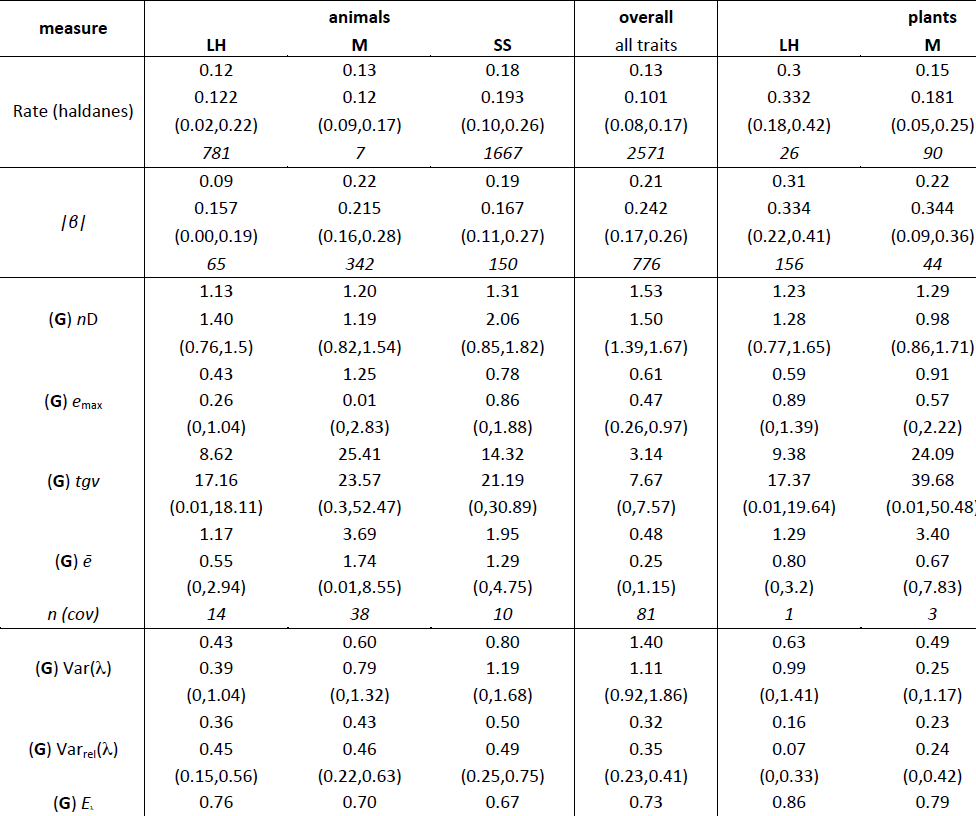
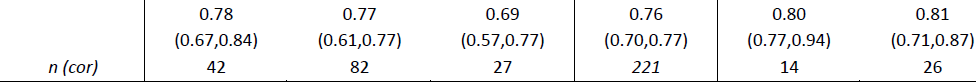
Summary statistics for estimates of the rate of evolutionary response, linear and quadratic selection gradients and measure capturing the size, shape and structure of **G**. (Statistics are reported by taxa and trait type, together with overall estimates across trait types and taxa. For each combination of taxa and trait type, the summary statistics for each measure are provided in the following order: posterior mean, posterior mode, lower and upper 95% credible intervals (in parenthesis) and sample size (in italics).)

## Results

### Observed rates of evolution differ among trait types and between plants and animals

The overall posterior mean for evolutionary rate was 0.13 haldanes, with a 95% credible interval from 0.08 − 0.17. Credible intervals for estimates in plants are quite wide (Figure 2), most likely due to the comparatively low number of studies in these categories. However there is a clear trend for faster rates in life-history traits, with the life-history estimate being ∼2.0 times as large (95% credible interval 0.7 − 4.8 x (the ratio calculated from MCMC iterations for both estimates)) as that for morphology, with only modest overlap of the 95% CI’s for the two trait types (Table 3). In animals, life-history and morphology have similar estimates, but the posterior mean estimate for sexually selected traits is somewhat higher – 1.5 times that for morphology (95% CI 0.5 − 6.9 times), and 1.5 times that for life-history (95% CI 0.8 − 2.3 times). Furthermore, the 95% CI’s for morphology do not include the estimate for sexually selected traits, though those for life-history do. Despite this, model support from various measures (AIC, BIC and DIC) is inconsistent about the overall support of trait types for the animal data improving model fit. Overall, these results suggest similar rates of evolution for morphology in both plants and animals, with higher rates for life-history traits in plants and possibly for sexually selected traits in animals.

**Figure 2.**
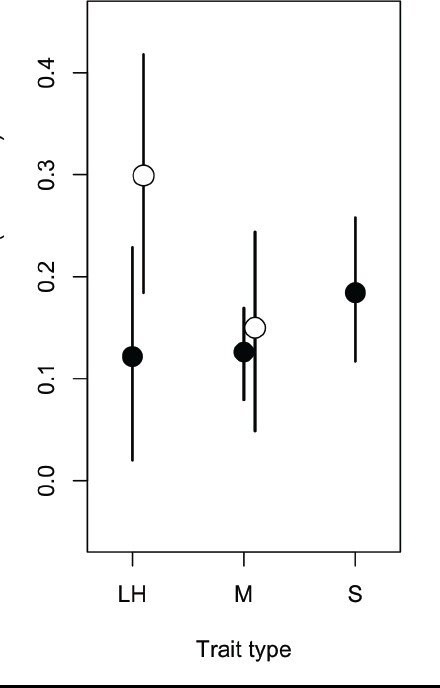
Posterior means and 95% credible intervals for estimates of absolute rate of evolution (haldanes). Open points are for plants and filled points for animals. Trait types are life-history (LH), morphology (M) and sexually selected (S) and filled points are for animals and open points for plants (no data available for sexual traits in plants).

### Standardised selection gradients show different patterns between plants and animals

The overall posterior mean for absolute linear selection gradients was 0.21 (95% CI = 0.17 − 0.26), which was somewhat higher than the estimate reported by Kingsolver et al. [38] (0.14, 95% CI = 0.13 − 0.16), most likely due to our inclusion of lab studies. The credible intervals from our full model are again wider for plants, likely reflecting smaller sample size (Table 3). For both plants and animals there is little difference between the estimates for morphological and sexually selected traits. In plants, the model suggests that selection is stronger on life-history traits, whose estimate is 40% larger than that for morphology and approximately twice that for sexually selected traits. By contrast, in animals selection appears to be weaker for life-history; the estimate for selection on life-history traits is 0.43 times (95% CI 0.11 − 0.97) that for morphology, and 0.49 times (95% CI 0.17 − 0.80) that for sexually selected traits (Figure 3).

**Figure 3.**
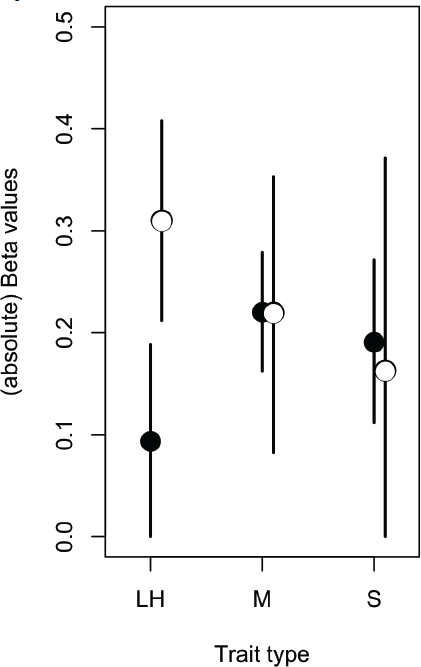
Posterior means and 95% credible intervals for estimates of standardized selection gradients (**β**) by trait type. Trait labels and taxon symbols are as in Figure 2.

### The marginal utility of multiple measures

The magnitude, shape and alignment of the **G** matrix all have the potential to influence the rate of evolution, but with the data available we are able to use measures intended to quantify only the first two of these properties. Of the measures (Table 1) we report *tgv*, *e*_max_ and *ē* can be thought of as measures of magnitude, whereas *n_D_*, Var(λ), Var_rel_(λ) and *E_λ_* are intended to quantify the departure of the matrix from symmetricality (how dissimilar variances are along the multiple axes of **G**). It is immediately obvious that the magnitude measures are doing a good job of quantifying the same property of each matrix (Table 1, Figures 4 & 5), since *tgv*, *e*_max_ and *ē* are all inter-correlated (*r* > 0.96 in all cases). Given that these measures of magnitude are also strongly correlated (*r* > 0.93 in all cases) with the magnitude of ***g****_max_* (i.e. the principal eigenvalue of **G**), it is perhaps unsurprising in retrospect that they are only poorly predicted by the number of traits measured, with which they are correlated only at *r* = 0.15 − 0.19.

**Figure 4.**
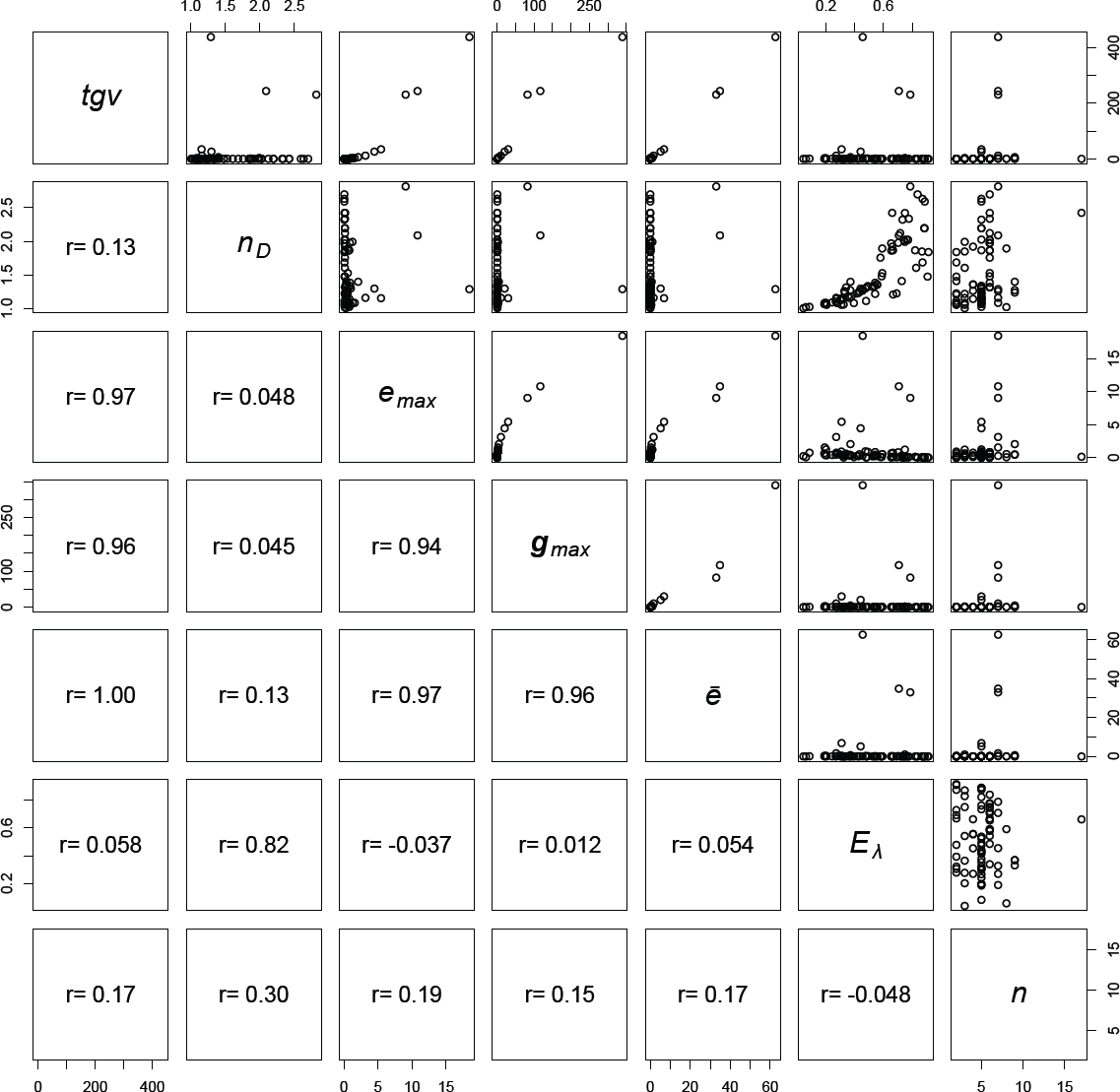
Pairs plot to illustrate the relationships between measures used to describe the structure of G expressed as covariance matrices. Measures are ‘total genetic variance’ (*tgv*), ‘maximum evolvability’ (*e*_max_) & ‘effective number of dimensions’ (*n_D_*) [20], the first eigenvalue of **G** (**g**_max_), ‘average evolvability’ (*ē*) [19], ‘eigenvalue evenness’ (*E_λ_* – originally intended for use with correlation matrices [23]) and the number of traits included in the matrix (*n*). Figures in the lower off-diagonal are pairwise correlations between the measures.

**Figure 5.**
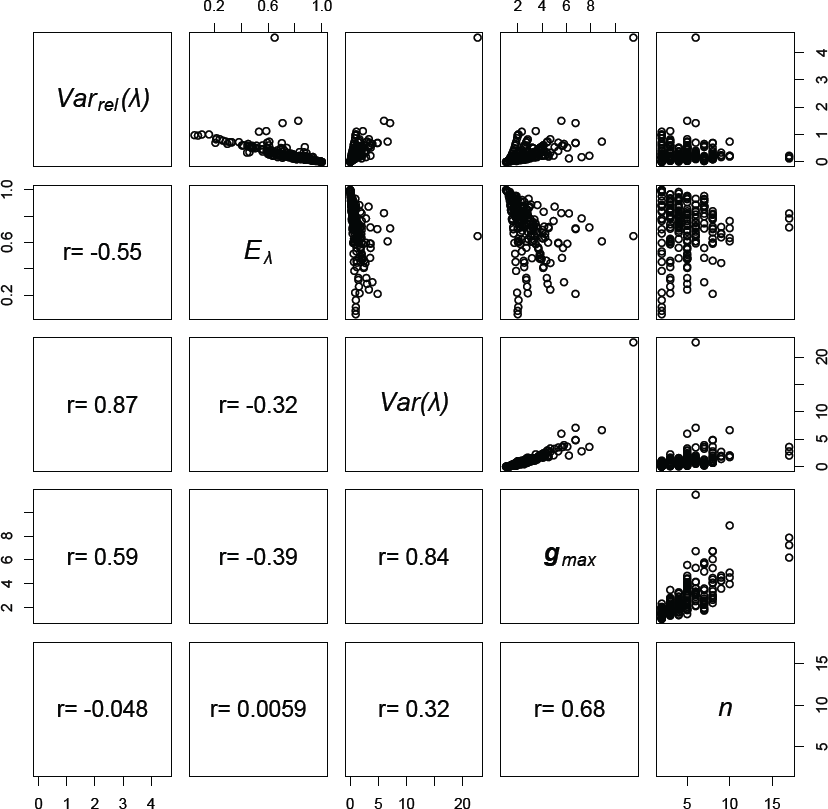
Pairs plot to illustrate the relationships between measures used to describe the structure of G expressed as correlation matrices. Measures are ‘relative eigenvalue variance’ (Var_rel_(λ)) [49], ‘eigenvalue evenness’ (*E_λ_*) [23], ‘eigenvalue variance’ (Var(λ)), [49], the first eigenvalue of **G** (**g**_max_) and the number of traits included in the matrix (*n*). Figures in the lower off-diagonal are pairwise correlations between the measures.

With respect to the measures of matrix eccentricity, the first thing we note is that Var(λ) and Var_rel_(λ) are strongly correlated with each other (*r* = 0.87), and negatively correlated with *E_λ_* (*r* = -0.32 & -0.55 respectively). Though *E_λ_* was defined as a measure of correlation matrices [23], when we applied the evenness formula to our dataset of covariance matrices we find that the resulting measure is strongly correlated with Kirkpatrick’s [20] *n_D_* (*r* = 0.82).

### The structure of **G**

We performed separate analyses and model selection procedures for each of our measures describing the structure of **G**. Our models comparing covariance matrices revealed very similar patterns of estimates for *e*_max_, *tgv* and *ē*. Furthermore the pattern of estimates among trait types was consistent between plants and animals (Figure 6). In all cases the estimates for life-history and sexually selected traits were similar and those for morphology were higher, but with much overlap in credible intervals our confidence in these differences is low. Our results for *n_D_* also show consistent patterns of estimates between plants and animals, with the estimates showing a shallow increasing trend from life-history to morphology to sexually selected traits (Figure 6(d)), but once again there is wide overlap among credible intervals, indicating low confidence in this trend. While this is for the inclusion of trait number as a linear covariate, similar results were obtained when *n_D_* was scaled directly by trait number (Figure S1).

**Figure 6.**
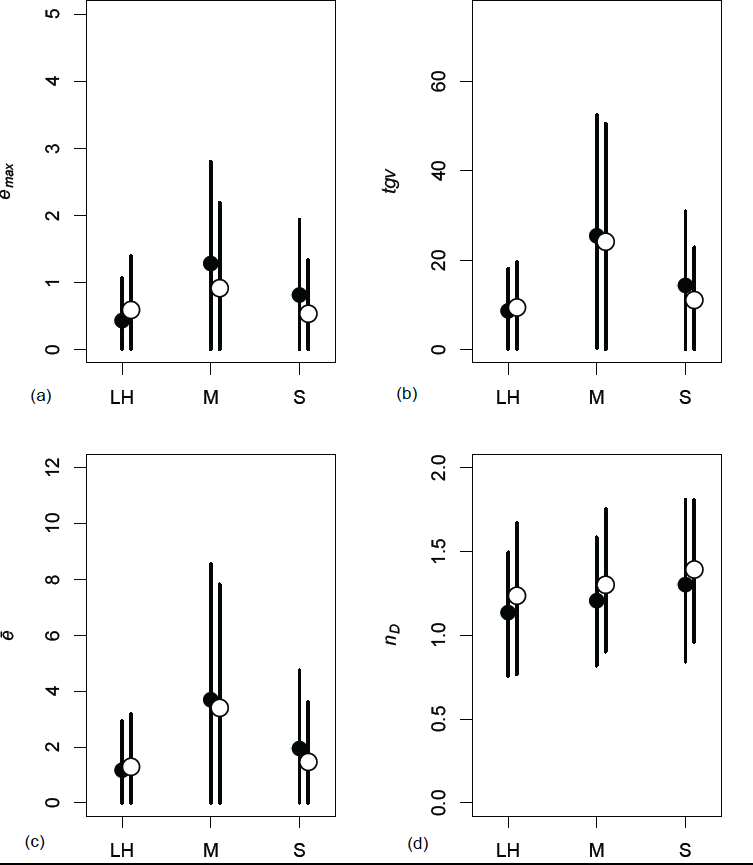
Posterior means and 95% credible intervals for the four measures used to characterise **G** matrices expressed as covariances (see methods section); (a) ‘maximum evolvability’ (*e*_max_), (b) ‘total genetic variance’ (*tgv*), (c) ‘average evolvability’ (*ē*) and (d) ‘effective dimensionality’ (*n_D_*). Trait types are life-history (LH), morphology (M) and sexually selected (S) and filled points are for animals and open points for plants.

The results of our analyses of **G** matrices expressed as correlations were more diverse. The pattern of estimates for Var_rel_(λ) showed a trend for values to increase from life-history to morphology to sexually selected traits in both plants and animals, though the estimates for animals were larger than those for plants (Figure 7(a)). The opposite trend was present in estimates for Var(λ) with the estimates for animals being somewhat lower than those for plants (Figure 7(b)). The wide overlap of credible intervals indicates low confidence in both these trends however. Finally, our estimates for *E_λ_* show a decreasing trend from life-history to morphology to sexually selected traits in both plants and animals, again with higher estimates for plants than for animals (Figure 7(c)).

## Discussion

**Figure 7.**
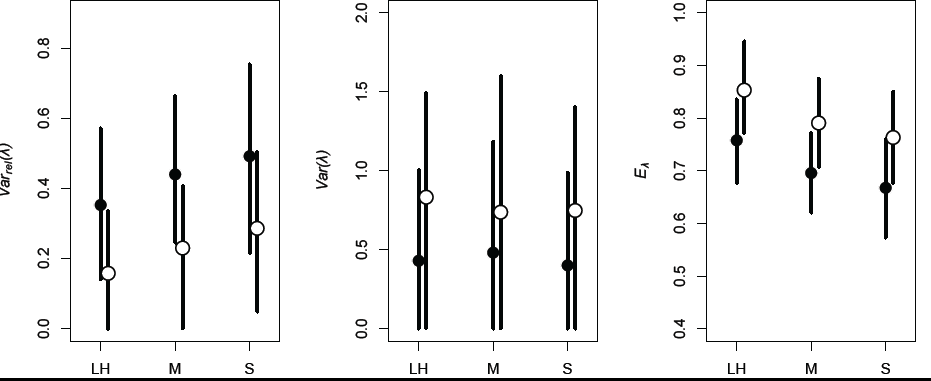
Posterior means and 95% credible intervals for the four measures used to characterise **G** matrices expressed as correlations (see methods section); (a) ‘relative eigenvalue variance’ (Var_rel_(λ)), (b) ‘eigenvalue variance’ (Var(λ)) and (c) ‘eigenvalue evenness’ (*E_λ_*). Trait labels and taxon symbols are as in Figure 6.

Predicting the rate and direction of phenotypic evolution is a fundamental challenge in evolutionary genetics [1–4,54], and the multivariate breeders’ equation is a key tool. Estimates of **G**, selection, and of response are available in the literature from many systems (though rarely reported together). Here we have integrated these data to ask if some traits evolve more rapidly than others, and whether differences associate with selection, **G** or both.

Reviews like this are unavoidably limited by the availability of published genetic parameters, and the resulting imbalances in the data. Nevertheless, we find some evidence that in animals – though not plants – sexual traits evolve faster than morphological traits. We find no evidence that this is due to stronger selection operating on these traits relative to morphological and life-history traits. We found weak evidence for differences in the evolutionary potential of **G** among trait types, though this fails to provide an explanation for any increased rates of evolution.

### Similarities among measures of the size and structure of **G**

We examined a number of the measures that have been proposed to assess the size, shape and structure of **G** (Table 1). Many of these measures have considerable shared information (Figures 4 & 5). Broadly, one group expresses the magnitude of **G** and a second relates to the evenness/variance of the eigenvalues, or eccentricity of **G**. While there may be particular instances where these measures result in widely divergent estimates, with respect to the empirical estimates we have collated, the marginal benefits of using all of them are an illustration of diminishing returns. It remains possible that subtle differences among these measures may provide important insights into the structure of **G** in the future. We speculate that one potential use (which would require considerable additional research) may be analogous to the population geneticists’ use of the parameter Tajima’s D, which is a scaled measure of two different estimates of the population mutation rate, 4N_e_µ.

One surprising observation that emerges from our results, is that the number of traits (*n*) used to estimate **G** is not well correlated with any of the measures we used. One potential explanation for this is that the magnitude of the principal eigenvalue of **G** is so highly correlated with ‘total genetic variation’ (the trace of **G**). This suggests that an overwhelming proportion of all of the variation is found along this principal vector (which would differ for each **G**), consistent with previous studies [9,20, 23]. It is known that estimating **G** can be difficult and insufficient sampling at the level of families can inflate the magnitude of the principal eigenvalue, at the expense of the minor eigenvalues [55,56]. However we saw no signal of such an effect from this database with any measures that capture eccentricity for **G** (Figures S4 & S5). As we did not have the raw data to re-compute **G** in a consistent framework, it is unclear how substantial this bias might be.

It is well known that scaling trait values by the mean versus the standard deviation can have profound impacts on univariate measures such as heritability. Likewise this would be expected for multivariate extensions like **G** and measures extracted from them as used here. Unfortunately in many instances the vector of trait means were unavailable, and thus our analysis for mean scaled **G** is a subset of that for the correlation matrices.

### Rates of evolution vary among traits

Reviews based on published estimates of evolutionary rates [30,31] have provided a number of important insights into the evolutionary process. Hendry & Kinnison [30] provided the foundations for measuring evolutionary rates and used a small sample of published estimates to propose that rapid evolution should be viewed as the norm rather than the exception. In a larger study, Kinnison & Hendry [31] showed that the frequency distribution of evolutionary rates measured in haldanes is log-normal (i.e. many slow rates and few fast rates, median haldanes = 5.8×10^-3^) and that life-history and morphological traits appear to evolve equally as fast when measured in haldanes. In agreement with these reviews, we found that the frequency distribution of evolutionary rates in our study was also log-normal and that the median rate across trait types and taxa was similar (median haldanes = 7.6×10^-3^) to that reported in Kinninson & Hendry [31]. We found little evidence to suggest that the evolutionary rates of life-history and morphological traits differed in animals, though there is evidence for faster rates in plant life-history. Our findings provide some evidence for a general pattern of faster evolution in sexual traits in animals to add to the highly cited individual examples of very rapid evolution of sexual traits [57,58] and their role in speciation [59,60]. It is worth noting that we used a different method for scaling data, as well as the inclusion of lab based studies of evolutionary rates, which differs from some other recent studies such as Uyeda et al. [46]. Future work examining how different methods of examining rate, and the inclusion of lab vs. field samples influence the overall observed pattern is warranted.

### The strength of selection varies among traits

Reviews synthesizing estimates of selection are extensive [33–39]. In their seminal review, Kingsolver *et al*. [33] found that the frequency distributions of linear and quadratic selection gradients were exponential and generally symmetrical around zero. This suggests that stabilizing and disruptive selection occur with equal frequency and with similar strength in nature. Kingsolver *et al.* [33] also found that the magnitude of linear selection was on average greater for morphological rather than life-history traits. The most recent review [38] containing an updated data set and using formal Bayesian meta-analysis to control for potential biases [34,35,37] confirmed many of the main findings of Kingsolver *et al*. [33], with the notable exception that linear selection appears stronger in plants than animals.

In agreement with this most recent synthesis [38], we found that the distribution of linear and quadratic selection gradients were exponential. Our estimates for absolute linear selection gradients were higher than reported by Kingsolver *et al.* [38] (0.24(0.17 − 0.26) versus 0.14 (0.13 − 0.16)). There has been much discussion on the general limitations of using selection gradients in synthetic reviews (e.g. [33,35,37,38]) and these arguments undoubtedly also apply to our study. However, as most of these limitations are inherent to both studies, they are unlikely to explain the observed differences. Furthermore, we used the same Bayesian framework as Kingsolver et al. [38] so it is unlikely that our analytical approach generated the observed differences. The most likely reason for the observed differences is the way that traits and taxa were categorized across these studies. Kingsolver et al. [38] used four different trait categories (size, morphological (not including size), phenology and life-history (not including phenology)) and categorized taxa as invertebrates, vertebrates or plants in their analysis. In contrast, we only distinguished between animals and plants and used three different trait categories (morphological, life-history and sexual) in our analysis, the latter of which includes a mixture of morphological and behavioural traits. Thus, there are likely to be some differences in how selection gradients are distributed amongst categories in our analyses compared to those in Kingsolver et al. [38]. Irrespective of the underlying reasons for these differences, we find little evidence for differences in the magnitude of selection gradients across trait types and taxa.

### Evolutionary response and the structure of **G**

After decades of quantitative genetic research it is now widely accepted that the additive genetic variance-covariance matrix (**G**) plays a major role in facilitating/constraining phenotypic evolution [16,19,20]. The way in which **G** shapes phenotypic evolution can be envisaged using the concept of genetic degrees of freedom (Figure 1; [9,15]). Whenever there is genetic covariation between the individual traits contained in **G**, there is the potential for fewer axes of genetic variation than observed traits [9,15,61,62] (but see [63]), which can influence evolutionary rates [64]. Where the majority of the genetic variance is concentrated in a few directions – known as “lines of least evolutionary resistance” (LLER’s) [15] – these have been shown to play an important role in directing the short-term evolutionary trajectory of a population [15,65–69]. Quantifying these properties of **G** is an essential step if we are to explore these ideas empirically. Perhaps unsurprisingly, it seems that the magnitude of a matrix is somewhat more straightforward to describe with a scalar measure than the eigenvalue evenness/eccentricity/dimensionality. The measures available for quantification of the shape of **G** in multiple dimensions are much less tightly inter-correlated than those dealing with matrix magnitude when compared using empirical data. What this ultimately means for our understanding of evolvability is unclear, but it is important to acknowledge the gaps in out current understanding if we are to progress.

Our finding that genetic variance for sexual traits may be spread less evenly across dimensions in animals runs counter to our hypothesis, and suggests that the potential for genetic constraint does not explain the higher rate of evolution we observe for these traits. We found at best, only weak evidence for differences in the measures to capture the size and shape of **G** with respect to our trait groupings. There has been debate over the importance of sexual selection in plants [70], but there is theoretical [48] and empirical [71] evidence suggesting that floral morphology is indeed subject to sexual selection. Unfortunately though, there are currently no data on evolutionary rates for sexual traits in plants, making it difficult to understand the implications of this increased dimensionality. Our findings indicate that the subject warrants greater attention.

### The effect of trait scaling

Researchers need to remain mindful that decisions about measurement scaling are likely to be important when measuring selection [35] and genetic variability [6]. This is especially important when addressing the question of evolvability, where both these measures must be brought together [19]. In this paper, we have attempted to present a clear picture of the patterns present in the currently available data, but it is important to acknowledge the known shortcomings of that data. This is not to understate the difficulty of maintaining comparability among studies wherein the appropriate scales might be different [6,35,72]. To illustrate the problem, how best to compare morphological data comprising linear measurements with life-history data where there may be no natural zero value? As a field, our inferences about selection and the response to selection will be more meaningful the more clearly we can address these issues.

### Conclusions

Collectively, our results suggest that the higher rate of evolution observed for sexual traits in animals is only weakly associated with the scalar measures summarizing **G** for these traits, and we do not find stronger selection. However, as our data set is based on derived estimates there are a number of inevitable limitations that apply to our findings. First, there are limitations with using the matrix structure measures (*n_D_*, *E_λ,_ Var*(*λ*) *or Var_rel_*(*λ*)) to capture the dimensionality of **G** [20]. Although these measure are calculable from published estimates of **G**, they do not explicitly test how many of the dimensions of **G** actually exist (i.e. have statistical support). A number of approaches [61,63] have been taken to directly estimate the dimensionality of **G** [61,73], though such studies have found both populations that have evolutionary access to all dimensions of **G** [63] and others that are constrained by LLER’s [61,74]. Second, our analysis does not consider the alignment between the vectors of selection and **G**. LLER’s only constrain the response to selection when they are poorly aligned with vectors of selection [26,28,64]. These limitations can only be resolved by further analysis of the raw data sets from the original studies we review. This is particularly true for better estimation of **G** itself, as well as its actual dimensionality, which can only be performed with the raw data [56,61,75–78]. Future studies would greatly benefit from researchers publishing raw datasets in open repositories [79] and we encourage researchers to do so. Our database (with all associated analyses) can be found at DRYAD DOI:xxxxxxxx, or on github (https://github.com/DworkinLab/Pitchers_PTRS2014).

## Acknowledgements

We thank S.F. McDaniel, R. L. Rodriguez and W.U. Blanckenhorn for contributing genetic data and to R. Snook, S. Chenoweth, T. Chapman, R. Firman, and M. Reuter for data on experimental evolution. I. Stott, F. C. Ingleby and three anonymous reviewers provided comments on earlier drafts of this manuscript, and A. Haber proofread the associated scripts. JH, TT and JBW were funded by NERC, JH and JBW by the BBSRC and JH by a University Royal Society Fellowship. WP was supported by a NERC studentship (awarded to JH and TT) and ID & WP were funded by NIH grant no. 1R01GM094424.

## Supplementary Material

### Further analyses of G matrix data

In response to a suggestion from reviewers, we model *n_D_* in 3 different ways. Initially, we had fitted the same suite of models that we used for the other G metrics, in addition to which we repeated the process with *n_D_/n* and also with *n_D_/n^2^*.

**Table S1:** The results of the model selection procedures for the 3 versions of *n_D_*.

| Response measure | Selected model |
| --- | --- |
| $n_D$ | trait type + taxon + trait no + random(study) |
| $n_D/n$ | trait type * taxon + random(study) |
| $n_D/n^2$ | trait type + taxon + random(study.code) + random(species) |

**Figure S1:**
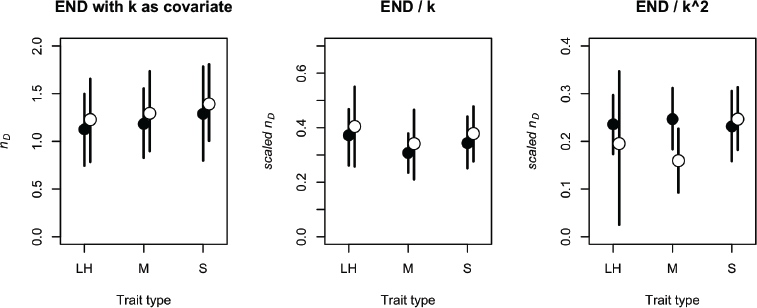
Results from alternative analyses of Kirkpatrick’s ‘effective number of dimensions’ metric.

**Figure S2:**
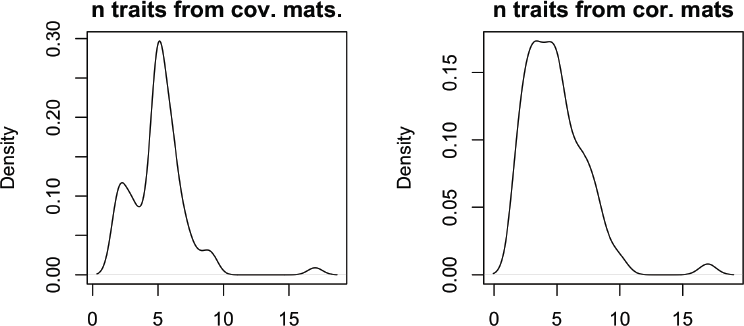
Density of the number of traits for out 2 G matrix datasets. Note that in both cases the majority of matrices are for between 4 & 6 traits. It is possible that there are effects associated with the number of traits that we have been unable to detect due to a lack of power. Only with a larger sample of larger matrices could we test this.

**Table S2:** The counts of numbers of matrices of each size (in terms of number of traits) represented in both our matrix datasets.

| $n$ | 2 | 3 | 4 | 5 | 6 | 7 | 8 | 9 | 10 | 17 | total |
| --- | --- | --- | --- | --- | --- | --- | --- | --- | --- | --- | --- |
| covariance matrix count | 11 | 8 | 5 | 29 | 16 | 6 | 2 | 3 | - | 1 | 81 |
| correlation matrix count | 31 | 41 | 36 | 42 | 21 | 21 | 17 | 5 | 4 | 3 | 221 |

We mentioned in the Discussion section that one legitimate concern with a quantitative review of the structure of **G** is that **G** can be challenging to estimate, and extremely challenging to estimate *well*. In particular, a smaller-than-optimal sample of families in a breeding design has the potential to inflate the magnitude of the **g**_max_, at the expense of the minor eigenvalues [55,56]. Given the importance of the ‘lines of least evolutionary resistance’ and ‘genetic degrees of freedom’ concepts for our thinking about multivariate evolution, it is a useful (not to mention reassuring) finding that there is no evidence to suggest that these patterns are driven by the sample sizes of the studies involved.

**Figure S3:**
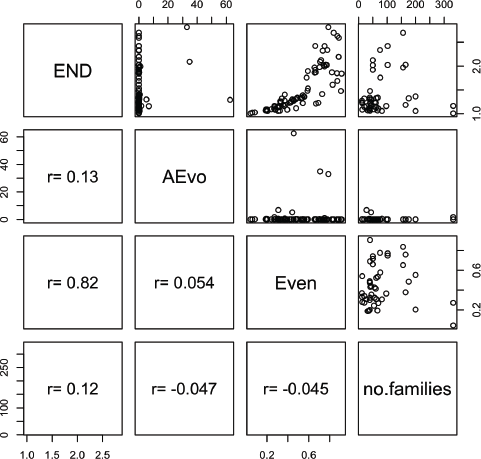
Pairs plot of the subset of covariance matrix measures that appear to represent the structure (as opposed to the magnitude) of **G**, in addition to the number of families measured to estimate **G**. (This plot does not include matrices estimated using an animal model, only those that result from breeding designs. ‘families’ in this case was taken to mean the number of sires in a half-sib design and the number of dams in parent-offspring regressions)

**Figure S4:**
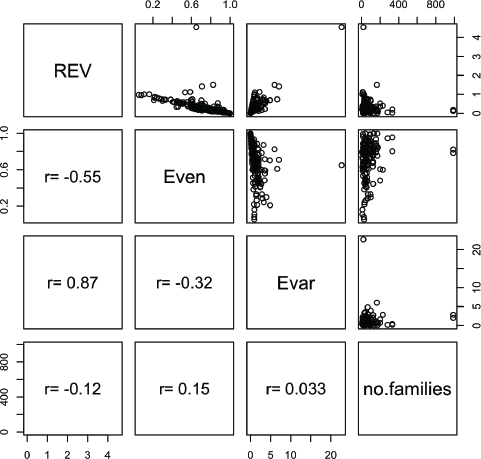
Pairs plot of the subset of correlation matrix measures that appear to represent the structure (as opposed to the magnitude) of **G**, in addition to the number of families measured to estimate **G**. (This plot does not include matrices estimated using an animal model, only those that result from breeding designs. ‘families’ in this case was taken to mean the number of sires in a half-sib design and the number of dams in parent-offspring regressions)

### Further analyses of selection data

In the main MS we only reported our findings from linear selection gradient (β) data. However, in the process of collecting these estimates we also tabulated estimates of quadratic selection gradients (the diagonal elements of the γ matrix). These estimates were reported less frequently than those for β, and there is a smaller dataset to work with. We divided the quadratic gradients into to groups; negative (potentially stabilizing) and positive (potentially disruptive) gradients. For each of these subsets, we fit the same model as used for the β dataset (a formal Bayesian meta-analysis following [38]: see main text), the results of which are visualized in Figure S5 below. Firstly, we should note that there are no differences among trait types or between taxa in which can have a high level of confidence. It is interesting to not that, for both taxa, there appear to be different trends in the two subsets of quadratic gradients, but we can say little more than that with the currently available data.

**Figure S5:**
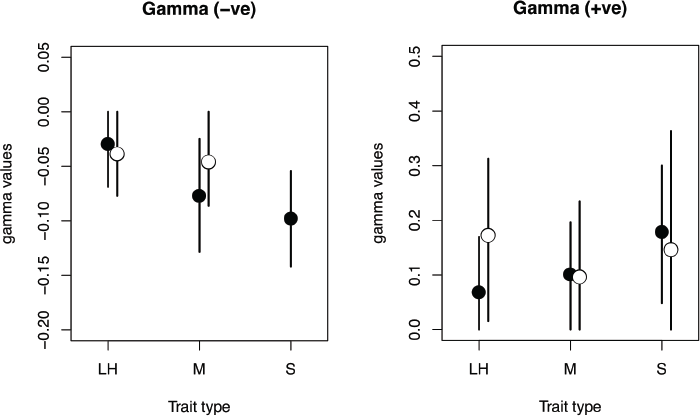
Posterior means and 95% credible intervals for negative (left panel) and positive (right panel) quadratic selection gradients. Trait types are life-history (LH), morphology (M) and sexually selected (S) and filled points are for animals and open points for plants.

## References

1. Arnold, S. J., Pfrender, M. E. & Jones, A. G. 2001 The adaptive landscape as a conceptual bridge between micro- and macroevolution. Genetica 112, 9–32.

2. Merila, J., Kruuk, L. E. B. & Sheldon, B. C. 2001 Natural selection on the genetical component of variance in body condition in a wild bird population. Journal of Evolutionary Biology 14, 918–929.

3. Begin, M. & Roff, D. A. 2003 The constancy of the G matrix through species divergence and the effects of quantitative genetic constraints on phenotypic evolution: A case study in crickets. Evolution 57, 1107–1120.

4. Walsh, B. & Blows, M. W. 2009 Abundant Genetic Variation + Strong Selection = Multivariate Genetic Constraints: A Geometric View of Adaptation. Annu. Rev. Ecol. Evol. Syst. 40, 41–59. (doi:10.1146/annurev.ecolsys.110308.120232)

5. Lande, R. 1979 Quantitative Genetic-Analysis of Multivariate Evolution, Applied to Brain - Body Size Allometry. Evolution 33, 402–416.

6. Houle, D. 1992 Comparing evolvability and variability of quantitative traits. Genetics 130, 195–204.

7. Roff, D. A. 1997 Evolutionary quantitative genetics. Springer.

8. Roff, D. A. & Mousseau, T. A. 2007 Quantitative genetics and fitness: lessons from *Drosophila*. Heredity 58 (Pt 1), 103–118. (doi:10.1038/hdy.1987.15)

9. Kirkpatrick, M. & Lofsvold, D. 1992 Measuring selection and constraint in the evolution of growth. Evolution 46, 954–971.

10. Bell, G. 1997 Selection: the mechanism of evolution. Springer.

11. Schluter, D. 2000 The Ecology of Adaptive Radiation (Oxford Series in Ecology & Evolution). 1st edn. Oxford University Press, USA.

12. Schluter, D. 2001 Ecology and the origin of species. Trends in Ecology & Evolution 16, 372–380.

13. Moose, S. P., Dudley, J. W. & Rocheford, T. R. 2004 Maize selection passes the century mark: a unique resource for 21st century genomics. Trends in Plant Science 9, 358–364. (doi:10.1016/j.tplants.2004.05.005)

14. Powell, R. L. & Norman, H. D. 2006 Major Advances in Genetic Evaluation Techniques. Journal of Dairy Science 89, 1337–1348. (doi:10.3168/jds.S0022-0302(06)72201-9)

15. Schluter, D. 1996 Adaptive radiation along genetic lines of least resistance. Evolution 50, 1766–1774.

16. Blows, M. W. & Hoffmann, A. A. 2005 A reassessment of genetic limits to evolutionary change. Ecology 86, 1371–1384.

17. Blows, M. W. 2007 A tale of two matrices: multivariate approaches in evolutionary biology. Journal of Evolutionary Biology 20, 1–8. (doi:10.1111/j.1420-9101.2006.01164.x)

18. Blows, M. & Walsh, B. 2009 Spherical cows grazing in flatland: constraints to selection and adaptation. 83–101.

19. Hansen, T. F. & Houle, D. 2008 Measuring and comparing evolvability and constraint in multivariate characters. Journal of Evolutionary Biology 21, 1201–1219. (doi:10.1111/j.1420-9101.2008.01573.x)

20. Kirkpatrick, M. 2008 Patterns of quantitative genetic variation in multiple dimensions. Genetica 136, 271–284. (doi:10.1007/s10709-008-9302-6)

21. Lande, R. & Arnold, S. J. 1983 The Measurement of Selection on Correlated Characters. Evolution 37, 1210–1226.

22. Hansen, T. F., Pélabon, C. & Houle, D. 2011 Heritability is not Evolvability. Evol Biol 38, 258–277. (doi:10.1007/s11692-011-9127-6)

23. Agrawal, A. F. & Stinchcombe, J. R. 2009 How much do genetic covariances alter the rate of adaptation? Proceedings of the Royal Society B: Biological Sciences 276, 1183–1191. (doi:10.1098/rspb.2008.1671)

24. Hunt, J., Wolf, J. B. & Moore, A. J. 2007 The biology of multivariate evolution. Journal of Evolutionary Biology 20, 24–27. (doi:10.1111/j.1420-9101.2006.01222.x)

25. Dickerson, G. E. 1955 Genetic Slippage in Response to Selection for Multiple Objectives. 20, 213–224. (doi:10.1101/SQB.1955.020.01.020)

26. Blows, M. W., Chenoweth, S. F. & Hine, E. 2004 Orientation of the Genetic Variance-Covariance Matrix and the Fitness Surface for Multiple Male Sexually Selected Traits. Am Nat 163, 329–340. (doi:10.1086/381941)

27. Hine, E., Chenoweth, S. F. & Blows, M. W. 2004 Multivariate quantitative genetics and the lek paradox: Genetic variance in male sexually selected traits of *Drosophila serrata* under field conditions. Evolution 58, 2754–2762.

28. Van Homrigh, A., Higgie, M., McGuigan, K. & Blows, M. W. 2007 The Depletion of Genetic Variance by Sexual Selection. Current Biology 17, 528–532. (doi:10.1016/j.cub.2007.01.055)

29. Simonsen, A. K. & Stinchcombe, J. R. 2010 Quantifying Evolutionary Genetic Constraints in the Ivyleaf Morning Glory, *Ipomoea hederacea*. Int. J Plant Sci. 171, 972–986. (doi:10.1086/656512)

30. Hendry, A. P. & Kinnison, M. T. 1999 Perspective: the pace of modern life: measuring rates of contemporary microevolution. Evolution 53, 1637–1653.

31. Kinnison, M. T. & Hendry, A. P. 2001 The pace of modern life II: from rates of contemporary microevolution to pattern and process. Genetica 112-113, 145–164.

32. Mousseau, T. A. & Roff, D. A. 1987 Natural selection and the heritability of fitness components. Heredity 59 (Pt 2), 181–197.

33. Kingsolver, J. G., Hoekstra, H. E., Hoekstra, J. M., Berrigan, D., Vignieri, S. N., Hill, C. E., Hoang, A., Gibert, P. & Beerli, P. 2001 The strength of phenotypic selection in natural populations. Am Nat 157, 245–261. (doi:10.1086/319193)

34. Hoekstra, H. E., Hoekstra, J. M., Berrigan, D., Vignieri, S. N., Hoang, A., Hill, C. E., Beerli, P. & Kingsolver, J. G. 2001 Strength and tempo of directional selection in the wild. Proceedings of the National Academy of Sciences 98, 9157–9160. (doi:10.1073/pnas.161281098)

35. Hereford, J., Hansen, T. F. & Houle, D. 2004 Comparing strengths of directional selection: how strong is strong? Evolution 58, 2133–2143.

36. Siepielski, A. M., DiBattista, J. D. & Carlson, S. M. 2009 It’s about time: the temporal dynamics of phenotypic selection in the wild. Ecology Letters 12, 1261–1276. (doi:10.1111/j.1461-0248.2009.01381.x)

37. Kingsolver, J. G. & Diamond, S. E. 2011 Phenotypic Selection in Natural Populations: What Limits Directional Selection? Am Nat 177, 346–357. (doi:10.1086/658341)

38. Kingsolver, J. G., Diamond, S. E., Siepielski, A. M. & Carlson, S. M. 2012 Synthetic analyses of phenotypic selection in natural populations: lessons, limitations and future directions. Evol Ecol 26, 1101–1118. (doi:10.1007/s10682-012-9563-5)

39. Siepielski, A. M., DiBattista, J. D., Evans, J. A. & Carlson, S. M. 2011 Differences in the temporal dynamics of phenotypic selection among fitness components in the wild. 278, 1572–1580. (doi:10.1098/rspb.2010.1973)

40. Morrissey, M. B. & Hadfield, J. D. 2011 Directional Selection in Temporally Replicated Studies Is Remarkably Consistent. Evolution 66, 435–442. (doi:10.1111/j.1558-5646.2011.01444.x)

41. Houle, D., Pélabon, C., Wagner, G. P. & Hansen, T. F. 2011 Measurement and meaning in biology. Q Rev Biol 86, 3–34.

42. Pomiankowski, A. & Møller, A. P. 1995 A resolution of the lek paradox. Proceedings of the Royal Society B: Biological Sciences 260, 21–29.

43. Haldane, J. B. S. 1949 Suggestions as to quantitative measurement of rates of evolution. Evolution 3, 51–56.

44. Gingerich, P. D. 1983 Rates of evolution: effects of time and temporal scaling. Science 222, 159–161. (doi:10.1126/science.222.4620.159)

45. Gingerich, P. D. 1993 Quantification and comparison of evolutionary rates. American Journal of Science 293-A, 453–478.

46. Uyeda, J. C., Hansen, T. F., Arnold, S. J. & Pienaar, J. 2011 The million-year wait for macroevolutionary bursts. P Natl Acad Sci Usa 108, 15908–15913. (doi:10.1073/pnas.1014503108)

47. Hendry, A. P., Farrugia, T. J. & Kinnison, M. T. 2008 Human influences on rates of phenotypic change in wild animal populations. Molecular Ecology 17, 20–29. (doi:10.1111/j.1365-294X.2007.03428.x)

48. Moore, J. C. & Pannell, J. R. 2011 Sexual selection in plants. Current Biology 21, R176–R182. (doi:10.1016/j.cub.2010.12.035)

49. Pavlicev, M., Cheverud, J. M. & Wagner, G. P. 2009 Measuring Morphological Integration Using Eigenvalue Variance. Evol Biol 36, 157–170. (doi:10.1007/s11692-008-9042-7)

50. R Core Dev. Team 2014. R: A language and environment for statistical computing.

51. Hadfield, J. D. 2010 MCMC methods for multi-response generalized linear mixed models: the MCMCglmm R package. Journal of Statistical Software 33, 1–22.

52. 2002 Bayesian measures of model complexity and fit. 64, 583–639.

53. Bates, D., Maechler, M., Bolker, B., & Walker, S. 2014. lme4: Linear mixed-effects models using Eigen and S4 (1st ed.) (http://CRAN.Rcproject.org/package=lme4)

54. Hill, W. G. & Zhang, X. S. 2012 On the Pleiotropic Structure of the Genotypec-Phenotype Map and the Evolvability of Complex Organisms. Genetics 190, 1131–1137. (doi:10.1534/genetics.111.135681)

55. Hill, W. G. & Thompson, R. 1978 Probabilities of Non-Positive Definite Between-Group or Genetic Covariance Matrices. 34, 429–439.

56. Meyer, K. & Kirkpatrick, M. 2010 Better Estimates of Genetic Covariance Matrices by ‘Bending’ Using Penalized Maximum Likelihood. Genetics 185, 1097–1110. (doi:10.1534/genetics.109.113381)

57. Mendelson, T. C. & Shaw, K. L. 2005 Sexual behaviour: rapid speciation in an arthropod. Nature 433, 375–376. (doi:10.1038/nature03320)

58. Bailey, N. W. & Zuk, M. 2008 Acoustic experience shapes female mate choice in field crickets. Proceedings of the Royal Society B: Biological Sciences 275, 2645–2650. (doi:10.1098/rsbl.2006.0539)

59. Panhuis, T. M., Butlin, R., Zuk, M. & Tregenza, T. 2001 Sexual selection and speciation. Trends in Ecology & Evolution 16, 364–371. (doi:10.1016/S0169-5347(01)02160-7)

60. Ritchie, M. G. 2007 Sexual Selection and Speciation. Annu. Rev. Ecol. Evol. Syst. 38, 79–102. (doi:10.1146/annurev.ecolsys.38.091206.095733)

61. Hine, E. & Blows, M. W. 2006 Determining the Effective Dimensionality of the Genetic Variance-Covariance Matrix. Genetics 173, 1135–1144. (doi:10.1534/genetics.105.054627)

62. McGuigan, K. & Blows, M. W. 2007 The Phenotypic and Genetic Covariance Structure of Drosphilid Wings. Evolution 61, 902–911. (doi:10.1111/j.1558-5646.2007.00078.x)

63. Mezey, J. G. & Houle, D. 2005 The dimensionality of genetic variation for wing shape in *Drosophila melanogaster*. Evolution 59, 1027–1038.

64. McGuigan, K. 2006 Studying phenotypic evolution using multivariate quantitative genetics. Molecular Ecology 15, 883–896. (doi:10.1111/j.1365-294X.2006.02809.x)

65. Badyaev, A. V. & Hill, G. E. 2000 The evolution of sexual dimorphism in the house finch. I. Population divergence in morphological covariance structure. Evolution 54, 1784–1794.

66. McGuigan, K., Chenoweth, S. F. & Blows, M. W. 2005 Phenotypic divergence along lines of genetic variance. Am Nat 165, 32–43.

67. Renaud, S., Auffray, J.-C. & Michaux, J. 2006 Conserved phenotypic variation patterns, evolution along lines of least resistance, and departure due to selection in fossil rodents. Evolution 60, 1701–1717.

68. Chenoweth, S. F. & McGuigan, K. 2010 The Genetic Basis of Sexually Selected Variation. Annu. Rev. Ecol. Evol. Syst. 41, 81–101. (doi:10.1146/annurev-ecolsys-102209-144657)

69. Boell, L. 2013 Lines of least resistance and genetic architecture of house mouse (*Mus musculus*) mandible shape. Evolution & Development 15, 197–204. (doi:10.1111/ede.12033)

70. Skogsmyr, I. O. & Lankinen, A. 2002 Sexual selection: an evolutionary force in plants? Biol. Rev. 77, 537–562. (doi:10.1017/S1464793102005973)

71. Carlson, J. E. 2008 Hummingbird responses to gender-biased nectar production: are nectar biases maintained by natural or sexual selection? Proc. Biol. Sci. 275, 1717–1726. (doi:10.1098/rspb.2008.0017)

72. Hansen, T. F., Pélabon, C., Armbruster, W. S. & Carlson, M. L. 2003 Evolvability and genetic constraint in Dalechampia blossoms: components of variance and measures of evolvability. Journal of Evolutionary Biology 16, 754–766. (doi:10.1046/j.1420-9101.2003.00556.x)

73. Mezey, J. G. 2005 Naturally Segregating Quantitative Trait Loci Affecting Wing Shape of *Drosophila melanogaster*. Genetics 169, 2101–2113. (doi:10.1534/genetics.104.036988)

74. Hunt, J., Blows, M. W., Zajitschek, F., Jennions, M. D. & Brooks, R. 2007 Reconciling Strong Stabilizing Selection with the Maintenance of Genetic Variation in a Natural Population of Black Field Crickets (*Teleogryllus commodus*). Genetics 177, 875–880. (doi:10.1534/genetics.107.077057)

75. Runcie, D. E. & Mukherjee, S. 2013 Dissecting high-dimensional phenotypes with bayesian sparse factor analysis of genetic covariance matrices. Genetics 194, 753–767. (doi:10.1534/genetics.113.151217)

76. de los Campos, G. & Gianola, D. 2007 Factor analysis models for structuring covariance matrices of additive genetic effects: a Bayesian implementation. Genet Sel Evol 39, 481–494. (doi:10.1051/gse:20070016)

77. Meyer, K. 2007 WOMBAT: a tool for mixed model analyses in quantitative genetics by restricted maximum likelihood (REML). J Zhejiang Univ Sci B 8, 815–821. (doi:10.1631/jzus.2007.B0815)

78. Mezey, J. G. & Houle, D. 2005 The dimensionality of genetic variation for wing shape in Drosophila melanogaster. Evolution 59, 1027–1038.

79. Whitlock, M. C., McPeek, M. A., Rausher, M. D., Rieseberg, L. & Moore, A. J. 2010 Data Archiving. Am Nat 175, 145–146. (doi:10.1086/650340)

